# LRTM: Left-Right Transition Matrices for Molecular Interaction Prediction

**DOI:** 10.1101/2025.02.06.636973

**Authors:** Kai Zheng, Guihua Duan, Mengyun Yang, Wei Wu, Yao-Hang Li, Jianxin Wang

## Abstract

Molecular interactions are central to most biological processes. The discovery and identification of potential associations between molecules can provide insights into biological exploration, diagnostic and therapeutic interventions, and drug development. So far many relevant computational methods have been proposed, but most of them are usually limited to specific domains and rely on complex preprocessing procedures, which restricts the models’ ability to be applied to other tasks. Therefore, it remains a challenge to explore a generalized approach to accurately predicting potential associations. In this study, We propose Left-Right Transition Matrices (LRTM) for molecular interaction prediction. From the perspective on the diffusion model, we construct two transition matrices to model undirected graph information propagation. This allows modeling the transition probabilities of links, which facilitates link prediction in molecular bipartite networks. The extensive experimental results show that the proposed LRTM algorithm performs better than the compared methods. Also, the proposed algorithm has the potential for cross-task prediction. Furthermore, case studies show that LRTM is a powerful tool that can be effectively applied to practical applications.

## 1 Introduction

Diseases are rarely caused by aberrations in a single gene, which reflect perturbations in complex intracellular and intercellular networks [1]. As a result, network medicine emerges as a necessity, because it not only identifies drug targets and biomarkers of complex diseases, but also reveals their biological significance and broadens our understanding of pathogenic mechanisms [2]. In network medicine, multiple molecular interactions are abstracted into bipartite networks, including disease-gene associations, drug-target interactions, drug-disease associations, lncRNA-diesease associations and miRNA-disease associations. These networks are all composed of two types of nodes and the edges that link them, and identifying interactions in networks can provide insights into biological exploration and therapeutic intervention [3]. Although some biotechnologies, e.g., proximity labeling [2], can discover endogenous interaction partners, they still suffer from high cost, long period of time, and labor-intensive. Consequently, computational methods are gaining traction as an economical alternative that, although not conclusive evidence, can provide valid candidates for biological experiments [4].

For the past several years, a number of computational methods have been proposed to identify potential associations for aiding clinical diagnosis and drug discovery, but most of them are designed for domain-specific tasks (e.g., drug repositioning, target identification, and disease-associated miRNA identification). These methods can be divided into two categories: the learning-based methods and the network-based methods. The learning-based methods usually characterize nodes as feature vectors and concatenate them together, which in turn transforms the link prediction problem into a binary classification problem. For example, Wang *et al*. predict disease-related miRNAs by using word2vec to extract miRNA sequence features and then combining multiple similarities to train the logistic model tree [5]. Zheng *et al*. proposed a chaos game representation-based miRNA representation method to extract miRNA sequence information, which improves the prediction performance of the broad learning system for miRNA-disease associations [6]. Guo *et al*. proposed a rotating forest-based model to predict disease-associated lncRNAs [7]. Shi *et al*. proposed an end-to-end model based on variational graph auto-encoders to identify potential lncRNA-disease associations. In addition to biomarker identification, drug-related association prediction is also a hot research topic [8]. Jiang *et al*. proposed a convolutional neural network based on the sigmoid kernel to predict the potential indications of drugs [9]. Zhao *et al*. enhanced the interpretability of drug-target binding affinity predictions by quantifying the importance of subsequences in an attention mechanism [10].

While the learning-based models achieve excellent performance, some of them are “opaque” to determine how they make decisions [11]. Also, they require large amounts of labeled data and multiple iterations to train their large number of parameters [12]. Recent studies show that the network-based approaches can identify antivirals that are not identified by the docking methods [13], [14]. Therefore, much research effort have focused on the networkbased methods. Thus far, many network-based models have been proposed to identify potential biological associations. For instance, Chen *et al*. proposed a computational model called MDHGI, which is based on the sparse learning method and the heterogeneous graph inference approach to identify miRNA-disease associations [15]. In order to accurately identify disease-related lncRNAs, Fu *et al*. proposed a computational method called MFLDA based on matrix tri-factorization, which reconstructs the association network by optimizing low-rank matrices [16]. Later, Wang *et al*. proposed a graph regularized non-negative matrix factorization-based model, which adopted K-nearest neighbors to reconstruct the lncRNA-disease association network [17]. In addition, drug target prediction and drug relocation are the research focus of network medicine. For example, Luo *et al*. proposed a method called MBiRW that integrated multiple similarities and utilized the Bi-Random Walk (BiRW) algorithm to identify new indications for drugs [18]. Recently, Ding *et al*. proposed a multi-kernel based triple cooperative matrix factorization (MK-TCMF) method to predict potential drug targets, in which multikernel learning was used to fuse multiple kinds of information [19]. Considering the fact that these methods rely on specific bipartite graphs and biological property information, generalization-capable, transferable, and interpretable methods are urgently needed.

We introduce Left-Right Transition Matrices (LRTM) as a universal approach for precise prediction of molecular interactions. By combining similarity and association networks, LRTM efficiently quantifies the significance of neighboring nodes surrounding a link, enabling accurate prediction of potential associations. Our model exhibits the following advantages: 1.LRTM serves as a versatile molecular interaction prediction model, adaptable to diverse scenarios. 2.The model provides transparency, offering a solid basis for inferring and interpreting prediction results with biological relevance. 3.LRTM models transition probabilities of links, facilitating assocition prediction in molecular bipartite networks. Extensive experimentation and evaluation demonstrate the effectiveness and potential utility of LRTM in predicting molecular interactions.

## 2 Materials and methods

### 2.1 Benchmark Datasets

To evaluate the generalization-capability and transferability of LRTM, we consider four types of biomedical bipartite networks, namely miRNA-disease association (MDA), lncRNA-disease association (LDA), drug-disease association (DDA) and drug-target Interaction (DTI). For DDA, we collect two public datasets, Fdataset [20] and Cdataset [18]. Here, we use two drug similarities. One is calculated by the semantic similarity algorithm [21] based on anatomical therapeutic chemical; the other is calculated by the Jaccard similarity coefficient based on the drug target network [22]. Also, two disease similarities are adopted. One is collected from MimMiner [23]; the other is calculated by using disease ontology terms. Details on disease and drug similarity can be found in the previous studies [24].

For DTI, we collect four publicly available datasets, namely enzymes (EN), nuclear receptors (NR), G-protein coupled receptors (GPCR) and ion channels (IC) [25]. Here, we use the drug similarity calculated by SIMCOMP [26] based on the chemical structure. The protein similarity is calculated by the Smith-Waterman score [27] based on the protein sequence.

For LDA, we adopt two public datasets, FuLDA and XieLDA [28]. In this study, we utilize the lncRNA similarity and the disease similarity provided by Fan *et al*. [28].

For MDA, we collect the dataset provided by Li *et al*. [29], HMDD v2.0. In detail, we obtain the miRNA similarity matrix by integrating the pre-calculated miRNA functional similarity and the Gaussian interaction profile kernel similarity matrix (GIP). The similarity between diseases comes from the pre-calculated disease semantic similarity and the GIP, where the semantic similarity is calculated by Medical Subject Headings (MeSH). The miRNA and disease similarities used are from the previous studies [30], [31]. It is worth noting that GIP is only based on the training set.

The detailed statistics of the nine biomedical bipartite networks for four types are shown in Table 1. It is worth noting that each association prediction problem involves many types of similarities, and similarities we used here are from comparison methods.

**TABLE 1.** AUC, AUPR and precision values of all compared methods in the ablation studies.

| Datasets | Type | Dimension | Edge | Density | Degree |
| --- | --- | --- | --- | --- | --- |
| Fdataset | DDA | (313, 593) | 1933 | 1.04% | 2.13 |
| Cdataset | DDA | (409, 663) | 2532 | 0.93% | 2.36 |
| EN | DTI | (664, 445) | 2926 | 0.99% | 2.64 |
| NR | DTI | (26, 54) | 90 | 6.41% | 1.23 |
| GPCR | DTI | (95, 223) | 635 | 3.00% | 2.00 |
| IC | DTI | (210, 240) | 1476 | 3.45% | 3.57 |
| HMDD | MDA | (495, 383) | 5430 | 2.86% | 6.18 |
| FuLDA | LDA | (240, 412) | 2697 | 2.73% | 4.14 |
| XieLDA | LDA | (115, 178) | 540 | 2.64% | 1.84 |

### 2.2 Construction of the heterogeneous network

In recent years, biological interaction networks have been constantly elucidated. Although we can simplify the processing procedure by modeling these molecules as homogeneous information networks, it is also prone to loss of information. Compared with homogeneous information networks, heterogeneous information networks naturally contain more molecular types and their interactions, thus containing richer semantics in nodes and links. Therefore, we utilize the heterogeneous information network to model complex relational data.

Based on previous works, we construct a heterogeneous network via the weighted similarity networks and the molecular bipartite network [24], [32]. The values of the similarities are between 0 and 1. We define the weighted similarities as **N1**_*S*_ and **N2**_*S*_, where N1 and N2 are the types of nodes.

By combining the weighted similarities **N1**_*S*_ and **N2**_*S*_ with the adjacency matrix **A**, we can define the bilayer network **A**_*h*_ as:

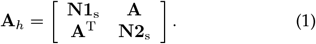

Here, taking the drug repositioning as an example, the construction process is shown in Fig. 1. We use the weighted average to fuse multiple similarities. The red and blue matrix in the Figure 1 represent the two drug similarities mentioned in Section 2.1. Similarly, the purple and brown matrix represent the two disease similarities.

**Fig. 1.**
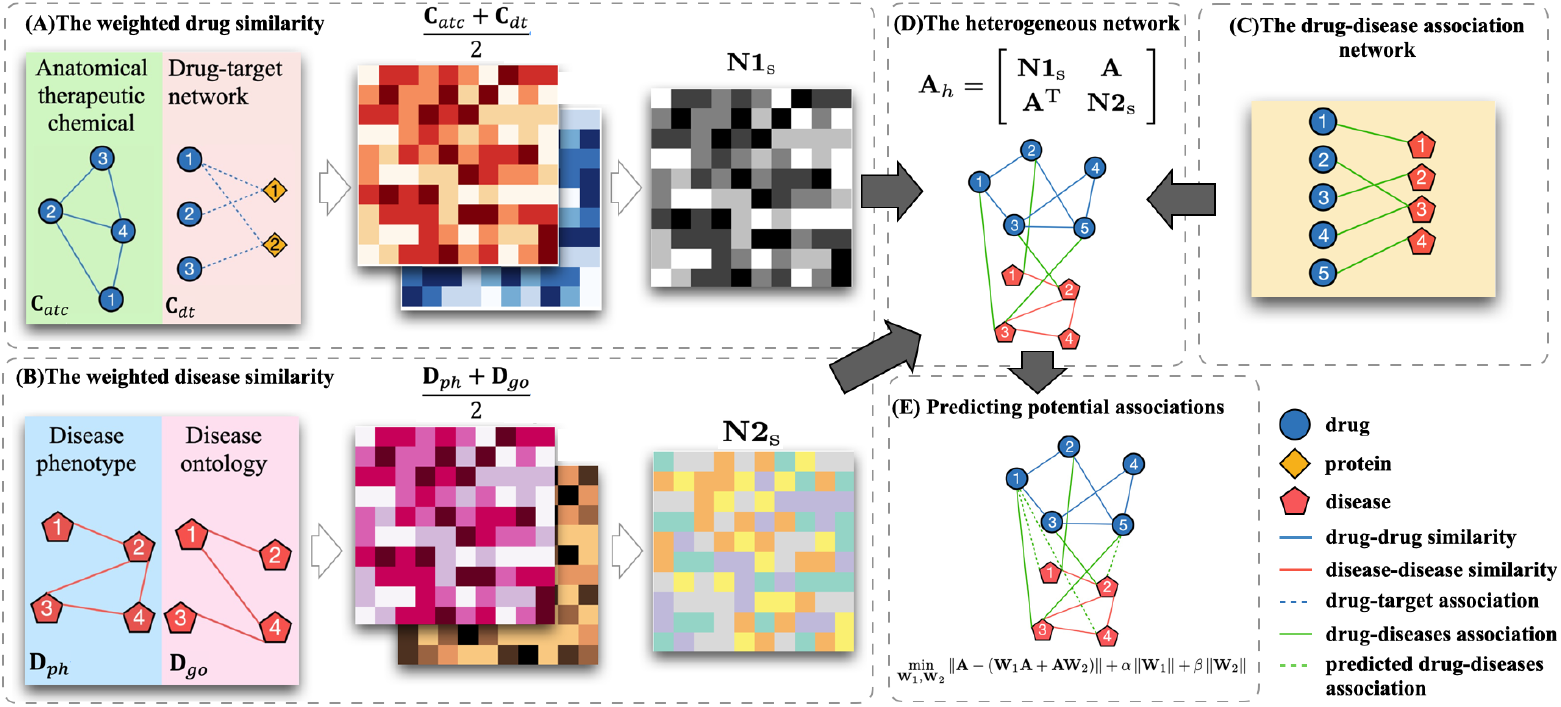
The workflow of the LRTM model for DDA. (A) The process of calculating weighted drug similarity. (B) The process of calculating weighted disease similarity. (C) The drug-disease association network. (D) The process of building the heterogeneous network. (E) Predict the potential associations by calculating **W**_1_, **W**_2_ in an optimization problem to determine the likelihood matrix **S**.

#### Algorithm 1

LRTM algorithm

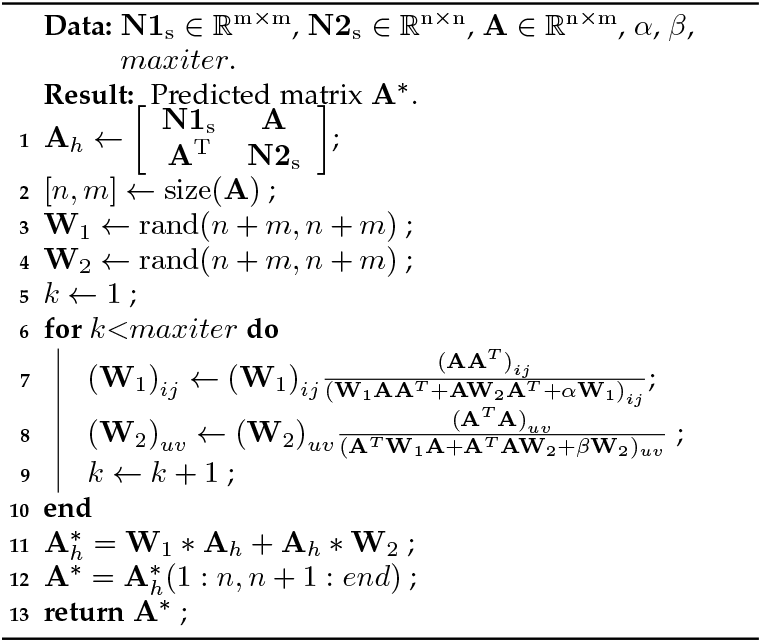

### 2.3 LRTM for molecular interaction prediction

Pech *et al*. proposed a link prediction method via linear optimization, LO [33], which rests on the assumption that the possibility of the existence of a link can be obtained by a linear summation of the contributions of neighboring nodes. The advantage of LO over other learning-based methods is that the weight matrix **W** can quantify the importance of each neighbor of a node, which provides valuable insights into the interpretability of the model(Fig. 2A). In addition, **W** can also be used as a graph node proximity metric for downstream task analysis and prediction. The link between node *i* and node *j* is considered only for the neighbors around node *i*. Because the model is designed for directed networks. However, when predicting links on undirected networks such as bipartite networks, the performance of this method may be degraded. For example, if *i* and *j* are different kinds of nodes, then the link between *i* and *j* is only relevant to nodes of the type to which *j* or *i* belong, and not to other type of node. Intuitively, the link between node *i* and node *j* should be related to the neighbors of both node *i* and node *j*. To improve the LO method, we propose a model called LRTM (as in Fig. 2B and Algorithm 1).

**Fig. 2.**
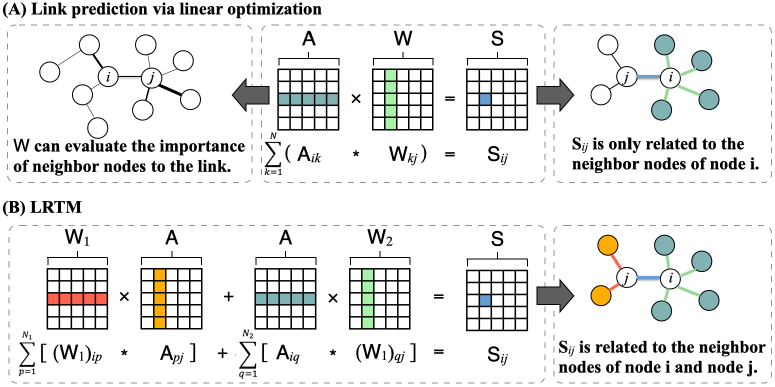
Comparison of link prediction via linear optimization and LRTM.(A) Link prediction via linear optimization. (B) LRTM.

We denote the molecular network as *G*(*V, E*), where *V* is the set of nodes and *E* is the set of edges. If node *i* and node *j* are associated, then **A**_*ij*_ = 1 in the corresponding adjacency matrix **A** ∈ ℝ^*m×n*^ (where *m* is the number of drugs and *n* is the number of diseases) of the molecular network, otherwise **A**_*ij*_ = 0. The likelihood matrix **S**_*ij*_ that *i* and *j* are associated can be defined as:

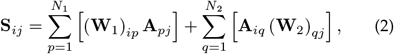

where (**W**_1_)_*ip*_ refers to the contribution of node *p* to node *j* in the association between *i* and *j*. According to the assumption, in the likelihood matrix, if **A**_*ij*¿_**A**_*pq*_, **S**_*ij*_ should also be larger than **S**_*pq*_. In other words, the difference between **A** and **S** should be small. In addition, to constrain **W**_1_ and **W**_2_ from becoming unitary matrices, we add two constraint terms to the optimization problem. Therefore, the likelihood matrix can be defined as an optimization problem:

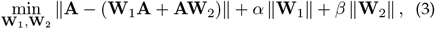

where *α* and *β* are the free parameters of the equilibrium optimization problem, and ∥ · ∥ is the matrix norm. Using the Frobenius norm with power 2, we transform the above equation into

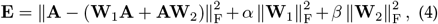

where 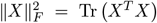. The expansion of the above formula reads

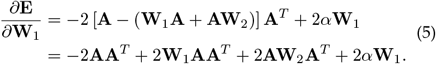

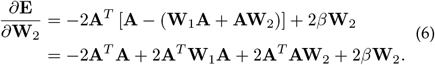

Setting ∂**E***/*∂**W**_1_ = 0 and ∂**E***/*∂**W**_2_ = 0, the weight matrices **W**_1_ and **W**_2_ can be expressed as:

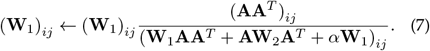

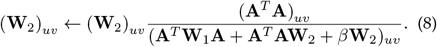

In summary, the novelty of the proposed method is that more neighboring nodes of the link are considered. From the perspective on the diffusion model, we construct two transition matrices to model undirected graph information propagation, which makes LRTM superior to LO in link prediction on the undirected networks in principle. In addition, LRTM inherits the advantage of LO, which quantifies the contribution of the neighboring nodes around the link. The contribution ratio *θ*_(*i,j*)_(*v*) of the neighbor node *v* of the link (*i, j*) can be expressed as follows:

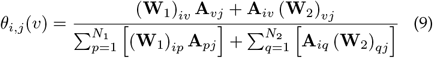

### 2.4 The time complexity analysis of LRTM

Matrix inversion and multiplication of three matrices are the two most computationally expensive parts of Eq.7 and Eq.8. The time complexity of matrix inversion is *O* (*s*^2.373^), where *s* is the size of the matrix [34]. The size of the adjacency matrix **A** is ℝ ^*m×n*^. In Eq.7, the time complexity of matrix inversion is *O* (*m*^2.373^), and the time complexity of **W**_1_ **AA**^T^ and **A W**_2_ **A**^T^ is *O* (m^2^*n*). In Eq.8, the time complexity of matrix inversion is *O*(*n*^2.373^), and the time complexity of **A**^T^**W**_1_**A** and **A**^T^**AW**_2_ is *O* (*n*^2^*m*). Therefore, we can get the time complexity of LRTM is *O* (max (*m*^2.373^, *m*^2^*n, n*^2.373^, *n*^2^*m*)).

## 3 RESULTS

### 3.1 Comparison with other baseline methods

To assess the predictive performance and generalization ability of the proposed method, a comparative analysis was conducted between LRTM and 10 state-of-the-art models using nine molecular bipartite networks (Table 2). Specifically, DRRS [35], OMC [36], HGIMC [32], and MSBMF [24] are employed for DDA prediction, L3 [37] and HGK [38] are utilized for DTI prediction, GCRFLDA [28] and DMFLDA [39] are employed for LDA prediction, and MDHGI [15] and NIMCGCN [40] are used for MDA prediction. We employed three evaluation metrics to assess the prediction performance: the area under the receiver operating characteristic (ROC) curve (AUC), the area under the Precision-Recall curve (AUPR), and Precision.

**TABLE 2.**
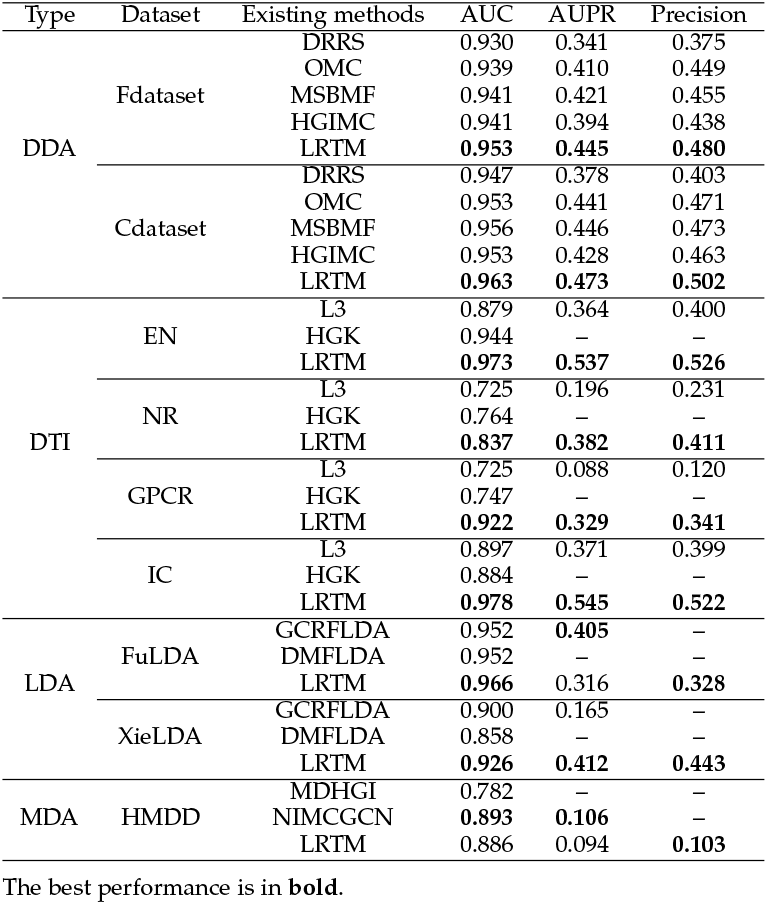
AUC, AUPR and precision values of all compared methods in nine molecular bipartite network.

**TABLE 3.** Performance comparison of DRRS, OMC, MSBMF, HGIMC and LRTM in *de novo* tests on Fdataset and Cdataset.

| Dataset | Metrics | DRRS | OMC | MSBMF | HGIMC | LRTM |
| --- | --- | --- | --- | --- | --- | --- |
| Fdataset | AUC | 0.824 | 0.851 | 0.875 | 0.877 | **0.901** |
|  | AUPR | 0.197 | 0.191 | 0.301 | 0.356 | **0.375** |
|  | Precision | 0.269 | 0.240 | 0.368 | 0.433 | **0.444** |
| Cdataset | AUC | 0.819 | 0.831 | 0.883 | 0.879 | **0.894** |
|  | AUPR | 0.193 | 0.170 | 0.298 | 0.294 | **0.329** |
|  | Precision | 0.254 | 0.226 | 0.367 | 0.356 | **0.401** |

The best performance is in **bold**.

For each of the compared models and datasets, we adhered to the recommended parameter settings and results as outlined in the respective publications. Table 2 presents a comprehensive performance comparison between LRTM and the related methods. It is worth noting that LRTM outperformed other methods on eight out of the nine datasets, showcasing its superior performance. Although LRTM was suboptimal for one dataset, its competitive performance across multiple datasets highlights its generalizability and transferability. Furthermore, we observe that LRTM performs well on tasks with low matrix density. For example, exceptional performance was achieved in Fdataset, Cdataset, and EN, where the density around 1% (refer to Table 1).

In addition, rigorous *de novo* tests were performed to evaluate the capability and efficacy of our method in predicting novel indications for drugs in DDA. This involved examining drugs with a sole known association, systematically removing each association to create a test sample, while utilizing the remaining associations as training samples [32]. By conducting these tests, we aim to provide compelling evidence of our method’s potential to predict previously unknown indications for drugs. These results suggest that LRTM exhibits great promise as an effective approach.

### 3.2 The sensitivity analysis of parameters

The proposed method involves two parameters, *α* and *β* of the equilibrium optimization problem in Equations 3 and 4. Possible values of *α* and *β* are 0 to +∞. Therefore, to demonstrate a reasonable search range for the parameters and a general trend of the impact on the prediction performance, we evaluate them by cross-validation. In order to evaluate the peak performance of all the algorithms and make the comparisons fair, we configure the key parameter settings of each algorithm on each data via grid search. Specifically, we select *α* and *β* from [0.1, 1, 10, 50, 100, 150, 300] in the case of the number of iterations, i.e., 30. Experiments are performed in nine biomedical networks. In this section, the average AUC is used as the metric to evaluate the hyperparameters.

Generally speaking, the optimal parameters of LRTM will be different on different benchmark datasets. As shown in Figure 3, in most cases, the optimal values of *α* and *β* lie in [1,50] except HMDD and fuLDA. Furthermore, we found that the parameter *β* had less effect on model performance in drug-related datasets. Consequently, we infer that drug similarity contributes more to the model’s decision-making, indicating that the current characterization of disease and protein similarity is still insufficient.

**Fig. 3.**
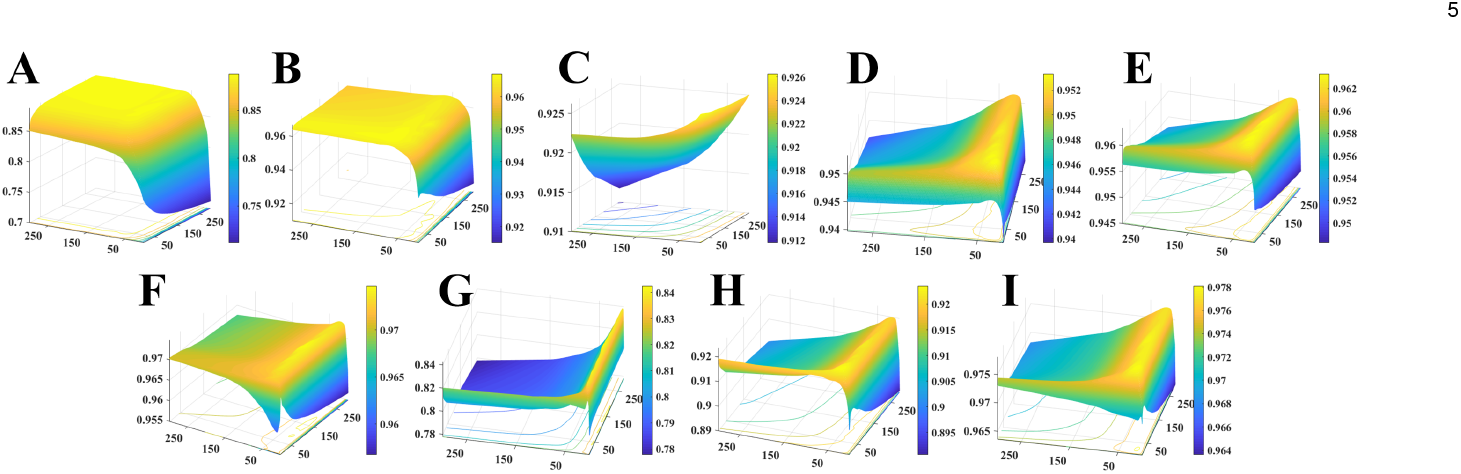
Performance of LRTM on nine molecular bipartite networks with different values of free parameters *α* and *β*: (A) HMDD, (B) FuLDA, (C) XieLDA, (D) Fdataset, (E) Cdataset, (F) EN, (G) NR, (H) GPCR, (I) IC.

**Fig. 4.**
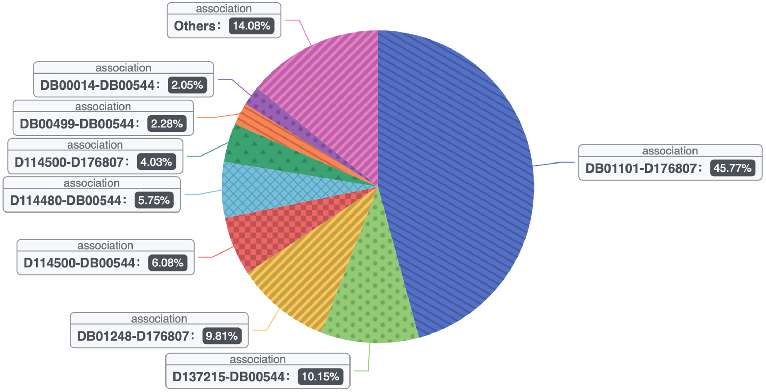
In the case study, LRTM’s decision basis for predicting prostate neoplasms (D176807) and fluorouracil (DB00544).

### 3.3 Comparison with LO

To compare the link prediction performance of LRTM and LO, we conducted two experiments in nine molecular bipartite networks, namely, the heterogeneous network **A**_*h*_ and the adjacency matrix **A** (section 2.2). The reason why we conduct the experiments with the adjacency matrices is that we can evaluate the raw performance of the model rather than assessing how well the model fits the similarity. In this section, the parameters of LO are selected from [0.0001, 0.001, 0.01, 0.1, 1]. The results show that the proposed method outperforms LO in most cases (Table 4), because the former introduces more neighbor nodes. In addition, the results also show that the link prediction performance is improved by constructing the heterogeneous network. It is worth noting that AUPR suffers from imbalanced datasets. Some methods select the same number of negative and positive samples. In contrast, all unlabeled samples were included in the calculations for this experiment.

**TABLE 4.**
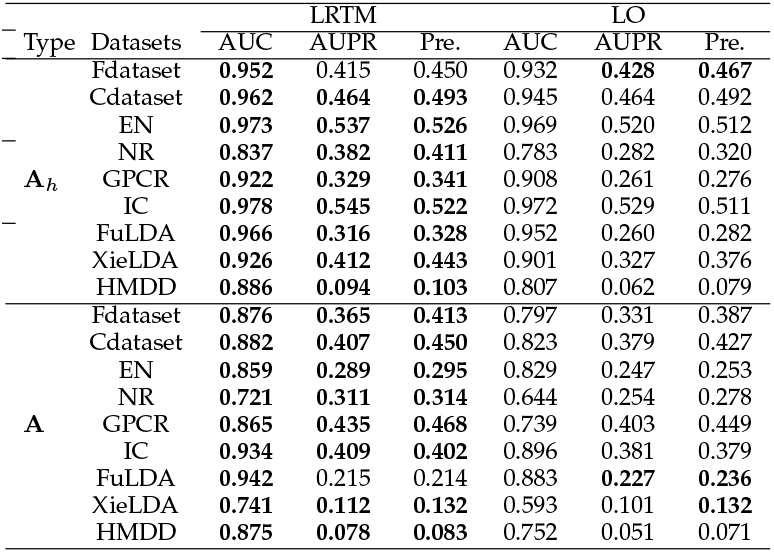
Performance comparison of LRTM and LO in nine molecular bipartite networks.

### 3.4 Case studies

To test the predictive performance of LRTM in reality, we identify the new indications for the approved drugs in the case studies. Specifically, Fdataset is used as a training set for scoring and ranking potential associations by LRTM. Here, four common drugs (Fluorouracil, Amantadine, Vincristine, and Methotrexate) are selected for case studies. As shown in Table 5, LRTM predicts the top ten indications with the highest scores for each of the four drugs. The indications that could be validated in the CTD database are bolded. For each representative drug, more than 3 candidates are reported in the public database, which demonstrates the potential of LRTM in realistically predicting new indications for drugs. In addition, we search the literature to validate the predicted top 10 drug-disease associations, 7 of which are confirmed (Table 6).

**TABLE 5.** Top 10 candidate indications for Fluorouracil, Amantadine, Vincristine and Methotrexate.

| Drugs (DrugBank IDs) | Top 10 candidate diseases (OMIM IDs) |
| --- | --- |
| Fluorouracil (DB00544) | **Prostatic Neoplasms (176807)**; Essential Hypertension (145500); **Urinary Bladder Neoplasms (109800)**; Osteosarcoma (259500); Leukemia, Acute Myelocytic, with Polyposis Coli and Colon Cancer (246470); **Esophageal Neoplasms (133239)**; Lymphoblastic Leukemia, Acute, with Lymphomatous Features (247640); Dohle Bodies and Leukemia (223350); Leukemia, Myeloid, Acute (601626); Hodgkin Disease (236000); |
| Amantadine (DB00915) | **Parkinson Disease (168600)**; **Dementia (125320)**; Restless Legs Syndrome (102300); Schizophrenia (181500); Insensitivity to Pain with Hyperplastic Myelinopathy (147530); Attention Deficit Disorder with Hyperactivity (143465); Sleep Apnea, Obstructive (107650); Hereditary spastic paralysis, infantile onset ascending (607225); Dystonia (601042); **Huntington Disease (143100)**; |
| Vincristine (DB00541) | Lymphocytic, Chronic, B-Cell (151400); **Osteosarcoma (259500)**; **Urinary Bladder Neoplasms (109800)**; **Lung Neoplasms (211980)**; Testicular Germ Cell Tumor (273300); **Breast Neoplasms (114480)**; Thyroid Cancer, Nonmedullary (188470); Small Cell Lung Carcinoma (182280); **Stomach Neoplasms (137215)**; **Mycosis Fungoides (254400)**; |
| Methotrexate (DB00563) | Neuroblastoma (256700); **Lung Neoplasms (211980)**; Wilms Tumor (194070); **Multiple Myeloma (254500)**; **Lymphocytic, Chronic, B-Cell (151400)**; Leukemia, Multicentric Carpotarsal Osteolysis Syndrome (166300); **Prostatic Neoplasms (176807)**; Purpura, Thrombocytopenic (188030); **Glioma (137800)**; **Lupus Erythematosus, Systemic (152700)**; |

The predicted indications in bold have been confirmed by CTD database.

**TABLE 6.** The top 10 drug-disease associations predicted by LRTM.

| Drug (DrugBank ID) | Disease (OMIM ID) | Evidence |
| --- | --- | --- |
| Enalapril (DB00584) | Renal Failure, Progressive, with Hypertension (161900) | [41] |
| Levetiracetam (DB01202) | Retinal Degeneration and Epilepsy (267740) | N/A |
| Levetiracetam (DB01202) | Ataxia with Myoclonic Epilepsy and Presenile Dementia (208700) | [42] |
| Diethylpropion (DB00937) | Chromosome Xq21 Deletion Syndrome (303110) | N/A |
| Zoledronic acid (DB00399) | Osteoporosis (166710) | [43] |
| Chlorothiazide (DB00880) | Ascites, Chylous (208300) | [44] |
| Metipranolol (DB01214) | Renal Failure, Progressive, with Hypertension (161900) | [45] |
| Ciclesonide (DB01410) | Asthma, Susceptibility To (600807) | [46] |
| Pamidronic acid (DB00282) | PAGET DISEASE OF BONE 2, EARLY-ONSET (602080) | [47] |
| Biperiden (DB00810) | Parkinson Disease, Late-Onset (168600) | N/A |

#### Interpretability

The contribution rate of each neighbor node of the link can be calculated by Equation 9, which enables the proposed model to provide a judgmental basis for the inference results. Taking prostate neoplasms (D176807) and fluorouracil (DB00544) as examples, we analyze whether the inference of the LRTM is in line with biological significance. We find that the three nodes with the highest contribution are all related to this association. Specifically, the neighbor node with the highest contribution is Capecitabine (DB01101, 45.8%), which slows the growth of neoplastic tissue by enzymatic conversion to fluorouracil [48]. Diffuse gastric and lobular breast cancer syndrome was the node with the second highest contribution (D137215, 10.1%), similar to prostate neoplasm, which is caused by mutations in the CDH1 gene [49], [50]. In the third place was docetaxel (DB01248, 9.8%), which, along with fluorouracil, can treat unresectable head and neck cancer with fluorouracil [51]. The results show that the proposed method is able to provide insights into mechanisms for prediction.

While numerous methods exhibit high accuracy, they often lack transparency, rendering them as “black boxes” that offer little insight into the underlying rationale. Consequently, the biological relevance of the inference process leading to their conclusions remains uncertain. In contrast, our proposed method not only achieves accuracy but also furnishes tangible evidence, enabling informed judgment regarding its biological significance.

## 4 Conclusion

In this study, we present a novel approach called Left-Right Transition Matrices (LRTM) for predicting molecular interactions. From the perspective on the diffusion model, we construct two transition matrices to capture the propagation of undirected graph information. This enables the modeling of transition probabilities for links, thereby facilitating link prediction in molecular bipartite networks. Through extensive experiments conducted on nine molecular interaction networks, LRTM consistently outperforms other models, underscoring its efficacy. Furthermore, case studies validate LRTM as a powerful and practical tool. However, there are still opportunities for enhancing our proposed model. Firstly, the computational efficiency is hindered by the involvement of matrix inversion, particularly in the context of large networks. Our future work will concentrate on reducing the time complexity associated with matrix inversion. Additionally, the performance of the proposed model can be influenced by the choice of similarity measures. Notably, the Jaccard similarity-based model outperforms the Pearson correlation coefficient-based model. Therefore, reducing the model’s reliance on similarities constitutes another avenue of exploration in our future research endeavors.

## Acknowledgment

Funding: This work has been supported by NSFC-Zhejiang Joint Fund for the Integration of Industrialization and Informatization (grant No. U1909208); the National Natural Science Foundation of China (No. 61772552, 61832019); 111 Project (No. B18059); Hunan Provincial Science and Technology Program (No. 2018WK4001); Scientific Research Fund of Hunan Provincial Education Department (No. 18B469); The Key Project of Hunan Education Department (NO.21A0478); The Natural Science Foundation of Hunan Province (NO.2022JJ30549); Fundamental Research Funds for the Central Universities of Central South University (2021zzts0206). This work was supported in part by the High Performance Computing Center of Central South University.

---

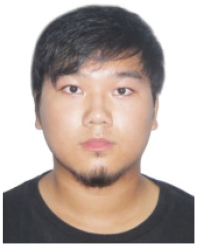

**Kai Zheng** received his B.E. degree in Computer Science and Technology from Central South University, Changsha, China, in 2017. He is currently a PhD student in Central South University. His current research interests include data mining, pattern recognition, deep learning, intelligent information processing and its applications in bioinformatics.

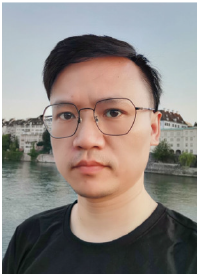

**Mengyun Yang** received the Ph.D. degree in computer science and technology from Central South University in 2020. He is an associate professor in the School of Science, Shaoyang University, Hunan, China. His current research interests include machine learning, deep learning, and bioinformatics.

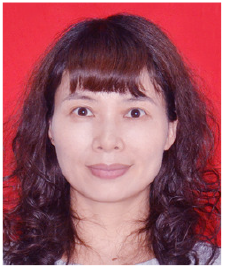

**Guihua Duan** received the PhD degree in computer science and technology from Central South University, in 2010. She is currently a professor with the School of Computer Science and engineering, Central South University. Her research interests include network security, bioinformatics, and computer network.

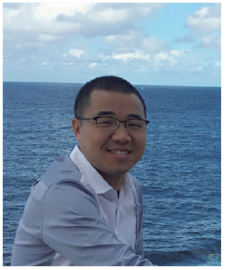

**Wei Wu** is an Associate Professor at School of Computer Science and Engineering, Central South University, China. His research interests are data mining and algorithms, and his papers appear in major conferences and journals including WWW, IJCAI, AAAI, ICDM, IEEE Transactions on Knowledge and Data Engineering and IEEE Transactions on Computers.

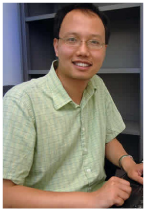

**Yaohang Li** received the BS degree from South China University of Technology in 1997, and the MS and PhD degrees in computer science from Florida State University, Tallahassee, FL, USA, in 2000 and 2003, respectively. He is an associate professor in computer science at Old Dominion University, Norfolk, VA, USA. His research interests are in protein structure modeling, computational biology, bioinformatics, Monte Carlo methods, big data algorithms, and parallel and distributive computing.

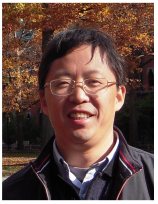

**Jianxin Wang** received the BEng and MEng degrees in computer engineering from Central South University, China, in 1992 and 1996, respectively, and the PhD degree in computer science from Central South University, China, in 2001. He is the dean and a professor in the School of Computer Science and Engineering, Central South University, Changsha, Hunan, China. He is a senior member of the IEEE. His current research interests include algorithm analysis and optimization, parameraized algorithm, bioinformatics, and computer network.

## References

[1] A.-L. Barabási, N. Gulbahce, and J. Loscalzo, “Network medicine: a network-based approach to human disease,” Nature reviews genetics, vol. 12, no. 1, pp. 56–68, 2011.

[2] W. Qin, K. F. Cho, P. E. Cavanagh, and A. Y. Ting, “Deciphering molecular interactions by proximity labeling,” Nature methods, vol. 18, no. 2, pp. 133–143, 2021.

[3] B. Munos, “Lessons from 60 years of pharmaceutical innovation,” Nature reviews Drug discovery, vol. 8, no. 12, pp. 959–968, 2009.

[4] N. Stephenson, E. Shane, J. Chase, J. Rowland, D. Ries, N. Justice, J. Zhang, L. Chan, and R. Cao, “Survey of machine learning techniques in drug discovery,” Current drug metabolism, vol. 20, no. 3, pp. 185–193, 2019.

[5] L. Wang, Z.-H. You, X. Chen, Y.-M. Li, Y.-N. Dong, L.-P. Li, and K. Zheng, “Lmtrda: Using logistic model tree to predict mirna-disease associations by fusing multi-source information of sequences and similarities,” PLoS computational biology, vol. 15, no. 3, p. e1006865, 2019.

[6] K. Zheng, Z.-H. You, L. Wang, Y.-R. Li, J.-R. Zhou, and H.-T. Zeng, “Missim: an incremental learning-based model with applications to the prediction of mirna-disease association,” IEEE/ACM Transactions on Computational Biology and Bioinformatics, vol. 18, no. 5, pp. 1733–1742, 2020.

[7] Z.-H. Guo, Z.-H. You, Y.-B. Wang, H.-C. Yi, and Z.-H. Chen, “A learning-based method for lncrna-disease association identification combing similarity information and rotation forest,” IScience, vol. 19, pp. 786–795, 2019.

[8] Z. Shi, H. Zhang, C. Jin, X. Quan, and Y. Yin, “A representation learning model based on variational inference and graph autoencoder for predicting lncrna-disease associations,” BMC bioinformatics, vol. 22, no. 1, pp. 1–20, 2021.

[9] H.-J. Jiang, Z.-H. You, K. Zheng, and Z.-H. Chen, “Predicting of drug-disease associations via sparse auto-encoder-based rotation forest,” in International Conference on Intelligent Computing. Springer, 2019, pp. 369–380.

[10] Q. Zhao, M. Yang, Z. Cheng, Y. Li, and J. Wang, “Biomedical data and deep learning computational models for predicting compound-protein relations,” IEEE/ACM transactions on computational biology and bioinformatics, vol. 19, no. 4, pp. 2092–2110, 2021.

[11] R. Shwartz-Ziv and N. Tishby, “Opening the black box of deep neural networks via information,” arXiv preprint arXiv:1703.00810, 2017.

[12] F. Sung, Y. Yang, L. Zhang, T. Xiang, P. H. Torr, and T. M. Hospedales, “Learning to compare: Relation network for few-shot learning,” in Proceedings of the IEEE conference on computer vision and pattern recognition, 2018, pp. 1199–1208.

[13] D. M. Gysi, Í. Do Valle, M. Zitnik, A. Ameli, X. Gan, O. Varol, S. D. Ghiassian, J. Patten, R. A. Davey, J. Loscalzo et al., “Network medicine framework for identifying drug-repurposing opportunities for covid-19,” Proceedings of the National Academy of Sciences, vol. 118, no. 19, 2021.

[14] G. Fiscon, F. Conte, L. Farina, and P. Paci, “Saverunner: a network-based algorithm for drug repurposing and its application to covid-19,” PLoS computational biology, vol. 17, no. 2, p. e1008686, 2021.

[15] X. Chen, J. Yin, J. Qu, and L. Huang, “Mdhgi: matrix decomposition and heterogeneous graph inference for mirna-disease association prediction,” PLoS computational biology, vol. 14, no. 8, p. e1006418, 2018.

[16] G. Fu, J. Wang, C. Domeniconi, and G. Yu, “Matrix factorization-based data fusion for the prediction of lncrna–disease associations,” Bioinformatics, vol. 34, no. 9, pp. 1529–1537, 2018.

[17] M.-N. Wang, Z.-H. You, L. Wang, L.-P. Li, and K. Zheng, “Ldgrnmf: Lncrna-disease associations prediction based on graph regularized non-negative matrix factorization,” Neurocomputing, vol. 424, pp. 236–245, 2021.

[18] H. Luo, J. Wang, M. Li, J. Luo, X. Peng, F.-X. Wu, and Y. Pan, “Drug repositioning based on comprehensive similarity measures and bi-random walk algorithm,” Bioinformatics, vol. 32, no. 17, pp. 2664–2671, 2016.

[19] Y. Ding, J. Tang, F. Guo, and Q. Zou, “Identification of drug–target interactions via multiple kernel-based triple collaborative matrix factorization,” Briefings in Bioinformatics, 2022.

[20] A. Gottlieb, G. Y. Stein, E. Ruppin, and R. Sharan, “Predict: a method for inferring novel drug indications with application to personalized medicine,” Molecular systems biology, vol. 7, no. 1, p. 496, 2011.

[21] P. Resnik, “Using information content to evaluate semantic similarity in a taxonomy,” arXiv preprint cmp-lg/9511007, 1995.

[22] P. Jaccard, “Nouvelles recherches sur la distribution florale,” Bull. Soc. Vaud. Sci. Nat., vol. 44, pp. 223–270, 1908.

[23] M. A. Van Driel, J. Bruggeman, G. Vriend, H. G. Brunner, and J. A. Leunissen, “A text-mining analysis of the human phenome,” European journal of human genetics, vol. 14, no. 5, pp. 535–542, 2006.

[24] M. Yang, G. Wu, Q. Zhao, Y. Li, and J. Wang, “Computational drug repositioning based on multi-similarities bilinear matrix factorization,” Briefings in Bioinformatics, vol. 22, no. 4, p. bbaa267, 2021.

[25] Y. Yamanishi, M. Araki, A. Gutteridge, W. Honda, and M. Kanehisa, “Prediction of drug–target interaction networks from the integration of chemical and genomic spaces,” Bioinformatics, vol. 24, no. 13, pp. i232–i240, 2008.

[26] M. Hattori, Y. Okuno, S. Goto, and M. Kanehisa, “Development of a chemical structure comparison method for integrated analysis of chemical and genomic information in the metabolic pathways,” Journal of the American Chemical Society, vol. 125, no. 39, pp. 11853–11865, 2003.

[27] T. F. Smith, M. S. Waterman et al., “Identification of common molecular subsequences,” Journal of molecular biology, vol. 147, no. 1, pp. 195–197, 1981.

[28] Y. Fan, M. Chen, and X. Pan, “Gcrflda: scoring lncrna-disease associations using graph convolution matrix completion with conditional random field,” Briefings in Bioinformatics, vol. 23, no. 1, p. bbab361, 2022.

[29] Y. Li, C. Qiu, J. Tu, B. Geng, J. Yang, T. Jiang, and Q. Cui, “Hmdd v2. 0: a database for experimentally supported human microrna and disease associations,” Nucleic acids research, vol. 42, no. D1, pp. D1070–D1074, 2014.

[30] K. Zheng, Z.-H. You, L. Wang, Y. Zhou, L.-P. Li, and Z.-W. Li, “Mlmda: a machine learning approach to predict and validate microrna–disease associations by integrating of heterogenous information sources,” Journal of translational medicine, vol. 17, no. 1, pp. 1–14, 2019.

[31] K. Zheng, Z.-H. You, L. Wang, Y. Zhou, L.-P. Li, and Z.-W. Li, “Dbmda: A unified embedding for sequence-based mirna similarity measure with applications to predict and validate mirna-disease associations,” Molecular Therapy-Nucleic Acids, vol. 19, pp. 602–611, 2020.

[32] M. Yang, L. Huang, Y. Xu, C. Lu, and J. Wang, “Heterogeneous graph inference with matrix completion for computational drug repositioning,” Bioinformatics, vol. 36, no. 22-23, pp. 5456–5464, 2020.

[33] R. Pech, D. Hao, Y.-L. Lee, Y. Yuan, and T. Zhou, “Link prediction via linear optimization,” Physica A: Statistical Mechanics and its Applications, vol. 528, p. 121319, 2019.

[34] X. Lei, X. Liao, T. Huang, H. Li, and C. Hu, “Outsourcing large matrix inversion computation to a public cloud,” IEEE Transactions on cloud computing, vol. 1, no. 1, pp. 1–1, 2013.

[35] H. Luo, M. Li, S. Wang, Q. Liu, Y. Li, and J. Wang, “Computational drug repositioning using low-rank matrix approximation and randomized algorithms,” Bioinformatics, vol. 34, no. 11, pp. 1904–1912, 2018.

[36] M. Yang, H. Luo, Y. Li, F.-X. Wu, and J. Wang, “Overlap matrix completion for predicting drug-associated indications,” PLoS computational biology, vol. 15, no. 12, p. e1007541, 2019.

[37] I. A. Kovács, K. Luck, K. Spirohn, Y. Wang, C. Pollis, S. Schlabach, W. Bian, D.-K. Kim, N. Kishore, T. Hao et al., “Network-based prediction of protein interactions,” Nature communications, vol. 10, no. 1, pp. 1–8, 2019.

[38] J. Lugo-Martinez, D. Zeiberg, T. Gaudelet, N. Malod-Dognin, N. Przulj, and P. Radivojac, “Classification in biological networks with hypergraphlet kernels,” Bioinformatics, vol. 37, no. 7, pp. 1000–1007, 2021.

[39] M. Zeng, C. Lu, Z. Fei, F. Wu, Y. Li, J. Wang, and M. Li, “Dmflda: a deep learning framework for predicting incrna–disease associations,” IEEE/ACM Transactions on Computational Biology and Bioinformatics, 2020.

[40] J. Li, S. Zhang, T. Liu, C. Ning, Z. Zhang, and W. Zhou, “Neural inductive matrix completion with graph convolutional networks for mirna-disease association prediction,” Bioinformatics, vol. 36, no. 8, pp. 2538–2546, 2020.

[41] A. Ciechanowicz, “Molecular mechanisms of nephro-protective action of enalapril in experimental chronic renal failure,” in Annales Academiae Medicae Stetinensis, 1999, pp. 1–93.

[42] C. R. Dolder and K. L. Nealy, “The efficacy and safety of newer anticonvulsants in patients with dementia,” Drugs & aging, vol. 29, no. 8, pp. 627–637, 2012.

[43] A. Räkel, A. Boucher, and L.-G. Ste-Marie, “Role of zoledronic acid in the prevention and treatment of osteoporosis,” Clinical Interventions in Aging, vol. 6, p. 89, 2011.

[44] S. Shaldon, J. Mclaren, and S. Sherlock, “Resistant ascites treated by combined diuretic therapy:(spironolactone, mannitol, and chlorothiazide),” The Lancet, vol. 275, no. 7125, pp. 609–613, 1960.

[45] J. Kuncová, J. Švíglerová, W. Kummer, D. Rajdl, M. Chottová-Dvořáková, Z. Tonar, L. Nalos, and M. Štengl, “Parasympathetic regulation of heart rate in rats after 5/6 nephrectomy is impaired despite functionally intact cardiac vagal innervation,” Nephrology Dialysis Transplantation, vol. 24, no. 8, pp. 2362–2370, 2009.

[46] R. S. Pirie, H.-W. Mueller, O. Engel, B. Albrecht, and M. von Salis-Soglio, “Inhaled ciclesonide is efficacious and well tolerated in the treatment of severe equine asthma in a large prospective european clinical trial,” Equine veterinary journal, vol. 53, no. 6, pp. 1094–1104, 2021.

[47] Z.-l. Zhang, X.-w. Meng, X.-p. Xing, O. Wang, W.-b. Xia, M. Li, Y. Jiang, Y. Fu, and X.-y. Zhou, “Prospective study of pamidronate disodium in treatment of paget’s disease of bone,” Zhonghua yi xue za zhi, vol. 83, no. 19, pp. 1653–1656, 2003.

[48] G. V. Koukourakis, V. Kouloulias, M. J. Koukourakis, G. A. Zacharias, H. Zabatis, and J. Kouvaris, “Efficacy of the oral fluorouracil pro-drug capecitabine in cancer treatment: a review,” Molecules, vol. 13, no. 8, pp. 1897–1922, 2008.

[49] P. R. Benusiglio, D. Malka, E. Rouleau, A. De Pauw, B. Buecher, C. Nogues, E. Fourme, C. Colas, F. Coulet, M. Warcoin et al., “Cdh1 germline mutations and the hereditary diffuse gastric and lobular breast cancer syndrome: a multicentre study,” Journal of medical genetics, vol. 50, no. 7, pp. 486–489, 2013.

[50] S. Lindström, F. Wiklund, B.-A. Jonsson, H.-O. Adami, K. Bälter, A. J. Brookes, J. Xu, S. L. Zheng, W. B. Isaacs, J. Adolfsson et al., “Comprehensive genetic evaluation of common e-cadherin sequence variants and prostate cancer risk: strong confirmation of functional promoter snp,” Human genetics, vol. 118, no. 3, pp. 339–347, 2005.

[51] J. B. Vermorken, E. Remenar, C. Van Herpen, T. Gorlia, R. Mesia, M. Degardin, J. S. Stewart, S. Jelic, J. Betka, J. H. Preiss et al., “Cisplatin, fluorouracil, and docetaxel in unresectable head and neck cancer,” New England Journal of Medicine, vol. 357, no. 17, pp. 1695–1704, 2007.

